# Mechanisms underlying the reprogramming of mouse embryonic fibroblasts to thymic epithelial cells

**DOI:** 10.1101/2022.03.21.485227

**Authors:** Zhongyao Ma, Seung Woo Kang, Brian G. Condie, Nancy R. Manley

**Affiliations:** Department of Genetics, University of Georgia, Athens, GA 30621

## Abstract

Thymic epithelial cells (TECs) are a critical functional component of the thymus’s ability to generate T cells for the adaptive immune system in vertebrates. However, no *in vitro* system for studying TEC function exists. Overexpression of the transcription factor FOXN1 initiates reprogramming of fibroblasts into TEC-like cells (iTECs) that support T cell differentiation in culture or after transplant. In this study, we characterized iTEC reprogramming at the cellular and molecular level to determine how reprogramming proceeds and identify mechanisms that can be targeted for improving this process. These data show that iTEC reprogramming consists of discrete gene expression changes that differ in early and late reprogramming, and that iTECs upregulate markers of both cortical and medullary TEC (cTEC and mTEC) lineages, although mTEC differentiation is blocked at a progenitor stage. We demonstrate that promoting proliferation enhances iTEC generation, and that Notch inhibition allows induction of mTEC differentiation. Finally, we show that a major difference between iTEC and fetal TEC is the expression of MHCII. This study supports future efforts to improve iTEC reprogramming for both research and translational uses.

## Introduction

The thymus is a primary lymphoid organ, and the major source of self-restricted, self-tolerant naïve T cells required for a robust adaptive immune system. However, through aging, the thymus is one of the earliest organs that starts losing its function, also known as thymic involution. Thymic involution leads to a decreased immune function, which significantly increases the risk of diseases, such as cancer and auto-immune diseases (Chinn et al., 2012; Hale et al., 2006; Palmer et al., 2018). Thus, finding an effective method to rescue thymus function caused by thymus involution and thymus abnormalities could have significant translational impact.

Thymic epithelial cells, or TECs, comprise the main functional resident cell types in the thymus. TEC-thymocyte interactions are required for T lineage commitment and all stages of thymocyte differentiation, proliferation, selection, and survival (Takahama, 2006). TECs are divided into cortical (cTEC) and medullary (mTEC) lineages, based on their location and function. There is some evidence that both lineages derive from a common progenitor, and that mTEC lineages derive from cells that have initiated cTEC differentiation (the ‘cTEC first’ model) (Baik et al., 2013; Bornstein et al., 2018; Rossi et al., 2006; Takahama et al., 2017). The mechanisms inducing cTEC differentiation are not understood. Recent evidence indicates that Notch signaling is required for mTEC lineage commitment but must be suppressed for mTEC differentiation to proceed past the progenitor stage (Li et al., 2020; Liu et al., 2020).

By E11.5, all cells specified for TEC identity within the 3rd pharyngeal pouch express the Forkhead transcription factor N1, Foxn1 (Vaidya et al., 2016). Null mutation of the *Foxn1* gene (nude) in mice, rats, and humans disrupts both normal hair growth and thymus development, resulting in immunodeficiency (Blackburn et al., 1996; Nehls et al., 1994). *Foxn1* is both necessary and sufficient for TEC differentiation and maturation and is the key transcription factor required for fetal TEC differentiation. FOXN1 also plays a critical role in postnatal thymus function and maintenance. *Foxn1* is widely expressed in cTECs and mTECs during postnatal stages, but is significantly decreased during aging, which is considered the major cause of thymic involution (Bredenkamp et al., 2014a; Chen et al., 2009; Ortman et al., 2002).

Although some evidence has shown that transplanting neonatal thymus or cytokine induction can partially rescue thymic function, these methods are all highly limited by resources and efficacy. Another approach is to use cellular reprogramming to generate key cell types *in vitro* for functional studies or transplantation. Direct lineage reprogramming is characterized by a dynamic and remarkable conversion of cellular morphology and transcriptomes. We and our collaborators have shown that over-expression of *Foxn1* is sufficient to reprogram mouse embryonic fibroblasts (MEFs) into functional induced thymic epithelial cells (iTECs). Within these *Foxn1+* primary MEFs, a subset not only began to express FOXN1 downstream targets *Dll4, Ccl25,* and *Kit-l,* but also Keratin-8 and epithelial cell adhesion molecule (EpCAM), which are expressed by all TECs during early thymus development. A subset of *Foxn1+* MEFs also started to exhibit epithelial cell-like morphology. Most importantly, iTECs could successfully support Early T cell progenitor (ETP) maturation into single positive (SP) T cells. Finally, iTECs grafted with supporting mesenchyme into a kidney capsule successfully developed into an organ with distinct cortical and medullary regions that could generate T-cells in the athymic nude mice. The result suggests the potential therapeutic application of iTECs. However, the process remains inefficient and has significant challenges to overcome before the *in vitro* system can be translational.

We used bulk RNA-seq to dissect the direct reprogramming process (Treutlein et al., 2016). We sequenced iTEC transcriptomes at different stages of reprogramming to characterize iTECs at the cellular and molecular level, which could identify potential mechanisms and pathways to improve the reprogramming. The results show that iTEC reprogramming consists of early and late reprogramming stages characterized by discrete gene expressions. In the early reprogramming stage, iTECs are characterized by activation of FOXN1 downstream targets, cellmatrix re-organization and cell morphology changes, and inhibition of cell cycle and cell proliferation. We demonstrate that releasing the proliferation block with an overexpression of Myc improves the reprogramming efficiency. In the late reprogramming stages, iTECs up-regulate additional epithelial markers and both cortical and medullary TEC (cTEC and mTEC) markers, although mature mTEC markers were not observed. We show that Notch signaling may block mTEC maturation, and that Notch inhibition induces the mTEC terminal differentiation marker *Aire.* Finally, we show that the main differences between iTECs and fetal TECs are the lack of expression of MHCII and the TEC-specific the transcription factor Tbata in iTECs, and an incomplete loss of fibroblast characteristics in iTECs. This study provides more detailed insights into the mechanisms behind the iTEC reprogramming by comparing transcriptomes between iTECs and fetal TECs and identifies several potential pathways to improve iTEC generation. Thus, this study provides new and critical information for developing iTECs as a useful experimental tool both for understanding *de novo* TEC biology and for potentially translational pre-clinical applications.

## Results

### Generation of iTECs using nucleofection based methods

To generate samples representing different iTECs reprogramming time-points, we first modified the originally published method of iTEC generation to use nucleofection based direct transfection of a *Cre*-expressing plasmid, instead of 4OHT treatment to activate an inducible CreER transgene as in the original iTEC protocol (Bredenkamp et al., 2014b). R26-CAG-Stop-Foxn1-IRES-GFP/+ (Fig.1A, B; R26-iFoxn1) heterozygous MEFs from E13.5 embryos were transfected with a PGK-Cre plasmid on day 0. After 48hrs of culture, a majority of the MEFs remove the STOP cassette by Cre recombinase and activate the R26-iFoxn1 locus, identified by GFP expression (Fig.1C). We termed these Foxn1- and GFP-expressing cells iFoxn1 MEFs. This method avoids the low level of “leaky” Cre expression from CreER, allowing us to precisely control the onset of iTEC reprogramming and providing a clean starting point for RNA-seq.

**Figure 1:**
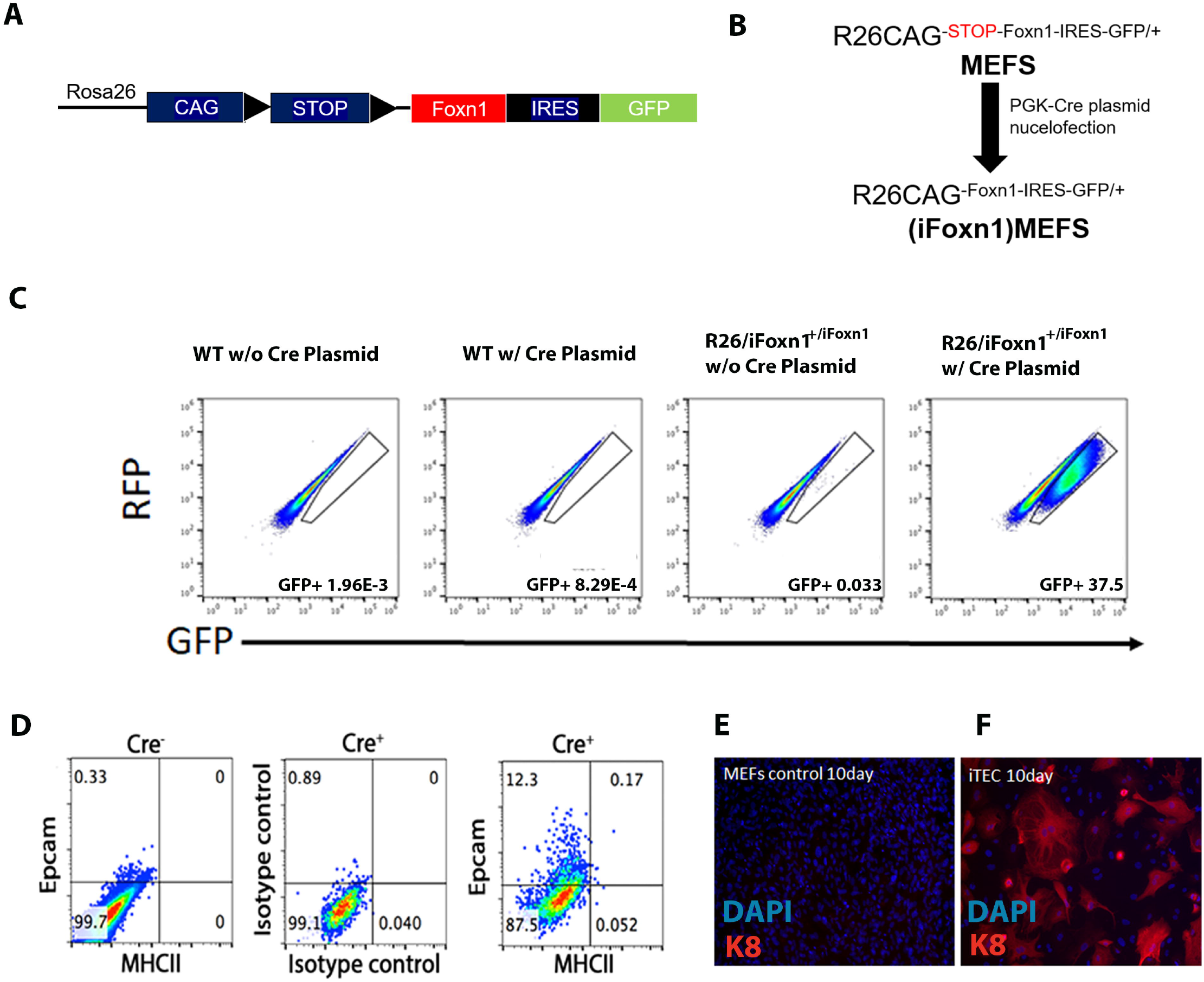
Generation and characteristics of R26^CAG-Foxn1-IRES-GFP/+^ MEFs. A) Gene structure of R26CAG-STOP-FOXN1-IRES-GFP transgene. B) Workflow of generation (iFoxn1) MEFS USING Cre-plasmid based nucleofection. C) IRES-GFP reporter signal analysis by flow cytometry for R26-ifoxn1/+ with Cre plasmid and other three control groups. D) Epithelium marker EpCam/MHCII analysis of R26-iFoxn1/+ and other 2 control groups with Cre+ plasmid 13 days after transfection. E) Epithelial marker Keratin 8 staining and DAPI staining of control MEFs. F) Keratin 8 staining and DAPI staining of iTEC 10day cells

As an initial characterization of the reprogramming process, we analyzed the iFoxn1 MEFs for the epithelium-specific marker EpCam and the TEC maturation marker MHC Class II (MHCII). Flow cytometry showed that more than 10% of iFoxn1 MEFs express EpCam 13days after the initial activation of transgenic *Foxn1* expression; however, as in the previous report, MHC Class II was not induced (Bredenkamp et al., 2014b) (Fig.1D). Also consistent with the previous report, many of the iFoxn1 MEFs were Keratin-8 positive 10 days after transfection (Fig.1F). No keratin-8 positive cells were observed in the mock transfected MEF controls (Fig. 1E).

We collected iFoxn1-MEFs at different time points after the initial iFoxn1 activation and used qRT-PCR to analyze the expression of several known FOXN1 target genes (Žuklys et al.) in TECs (Fig. 2A, B). FOXN1 target genes were detected early after induction of *Foxn1* expression, with *Dll4* and *Kit-L* significantly increased as early as 24hrs after transfection, and *Ccl25* is significantly up-regulated 2 days (2d) after transfection (Fig. 2A). The increase in these FOXN1 target genes over the course of the 12d iTEC induction period does not follow a linear pattern. Instead, these three genes undergo rapid expression increases starting at 7days after transfection, accelerating further after 10d (Fig. 2B). This pattern suggests that reprogramming has two different stages, an early stage with initial low-level expression of target genes, followed by a later stage in which FOXN1 target gene expression is dramatically accelerated.

**Figure 2:**
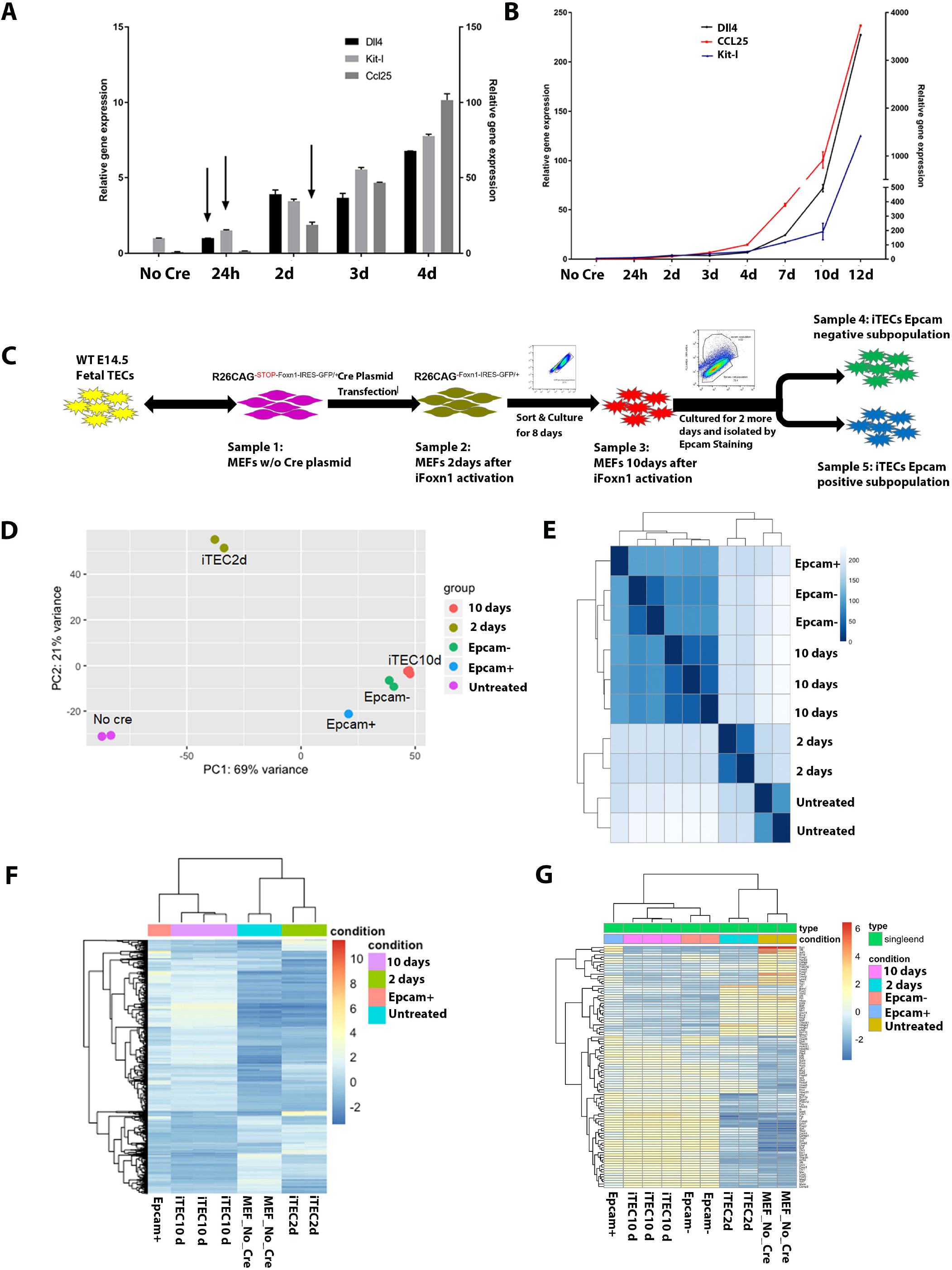
Schematic diagram of RNA-seq experiment design and general transcriptome analysis. A) Gene expression analysis od selected Foxn1 target genes using qRT-PCR at different timepoints after Cre transfection from 24 hrs to 4 days. B) Gene expression analysis using qRT-PCR at different time-point after Cre transfection from 24 hrs to 12 days. C) Schematic diagram and sample FACS sorting plot from generating iTEC samples for RNA-seq analysis. D) PCA analysis of all sample transcriptomes. E) Sample distance analysis of all sample transcriptomes. F) Top differentially expressed genes heatmap analysis of all samples. G) Top differentially expressed transcription factor heatmap analysis.

### iTEC reprogramming has progressive gene expression changes with a distinct intermediate cell step

To further understand the dynamic process of iTEC reprogramming, we designed a timepoint based bulk-RNA-seq experiment, choosing our time points based on the induction of FOXN1 target genes (Fig. 2C). We used mock-transfected R26-iFoxn1/+ MEFs as our control, representing the initial MEFs transcriptome status (No Cre MEFs). We induced R26-iFoxn1/+ MEFs to express *Foxn1* using Cre transfection, cultured for 2days, and sorted cells that had successfully induced *iFoxn1* expression as GFP+ using flow cytometry (as in Fig. 1C). These GFP+ iFoxn1 MEFs represent initial reprogramming and are referred to as iTEC2d. We also cultured iTECs for a total of 10days after transfection to represent the late stage of reprogramming, termed iTEC10d. For the final stage, we cultured iTECs for a total of 12 days and then used Epcam antibody staining to separate them into two distinct groups, designated EpCam+ & EpCam-. Each RNA-seq sample consisted of cells pooled from multiple independent iTEC cultures, with RNA-seq performed in duplicate (MEFs, iTEC2d, and EpCam-) or triplicate (iTEC10d). The exception is the EpCam+ iTECs, for which only one pooled sample was analyzed in these initial experiments.

To get an initial understanding of the progression of iTEC reprogramming, we first analyzed MEF and all iTEC whole transcriptome profiles using PCA analysis (Fig. 2D). We found that even 48hrs after the initial activation of ectopic Foxn1, iTEC gene expression has changed significantly compared to MEFs as indicated by the clear separation of iTEC2d and No Cre MEF samples. The iTEC10d sample is further separated from the iTEC 2d sample but does not show a linear increase of variance from No Cre to iTEC10d through iTEC2d, indicating that the iTECs reprogramming process is not a continuous process at the gene expression profile level but includes distinct gene expression pattern changes at the early and late stages. Finally, the EpCam+ samples are more distinct than EpCam-samples compared to iTEC 10day. The whole transcriptome analysis shows a linear progression of differential gene expression, which indicating that EpCam+ cells represent a more advanced reprogramming state, and suggesting that 10d, EpCam-, and EpCam+ iTEC represent a linear progression of differentiation (Fig. 2D). The sample distance count matrix also shows this progression process from MEFs to EpCam+ iTECs, with a significant difference between each step of reprogramming (Fig. 2E). Furthermore, the reproducibility of the samples at each reprogramming stage is high, with the exception of the EpCam+ stage, for which only one sample was analyzed.

The gene count matrix results show that each stage of reprogramming has a distinct signature, with sets of genes turning off and on across the reprogramming timeline. While there was a block of genes that were downregulated from MEFs to 2d then stayed low throughout reprogramming, relatively few genes showed a gradual upregulation from MEFs through the EpCam+ stage. For example, distinct subsets of genes are transiently expressed at both the iTEC2d stage (off in MEFs, on in iTEC2d, and off in iTECs10d and EpCam+) and the iTEC10d stage (off in MEFs and 2d, high in iTEC10d, then down again in EpCam+ iTECs) (Fig.2F). We specifically assessed transcription factor expression changes during the reprogramming process, comparing MEFs, 2d, 10d, and EpCam+ (Fig. 2G). Similar to the global DEG analysis, we find that specific sets of transcription factors were differentially expressed at the early and late reprogramming stages.

Finally, we separately analyzed up-regulated & down-regulated genes at different reprogramming stages by gene enrichment analysis (Fig. 3). We find that in early reprogramming stages from MEFs to iTEC2d, iTECs are characterized by down-regulation of cell shape, extracellular matrix (ECM), and other fibroblast related genes, and up-regulation of cytokine signaling and other immune-related categories (Fig. 3A, B). Between 2d and 10d, the cell cycle and DNA replication are inhibited, and genes related to epithelial characteristics are upregulated, indicating that the iTECs are transitioning into epithelial cells during this second stage of reprogramming (Fig. 3C, D). The distinctions between EpCam- and EpCam+ 12d iTECs are also made more clear when each are compared to iTEC10d. EpCam-iTECs upregulate programmed cell death and cell cycle inhibition categories, while down regulating epithelial-related genes, suggesting that they constitute cells that fail to maintain the transdifferentiation process and may undergo cell death (Fig. 3E-F). In contrast, EpCam+ cells upregulated differentiation and metabolism related gene categories, while downregulating categories related to mesenchymal properties (Fig. 3G-H).

**Figure 3:**
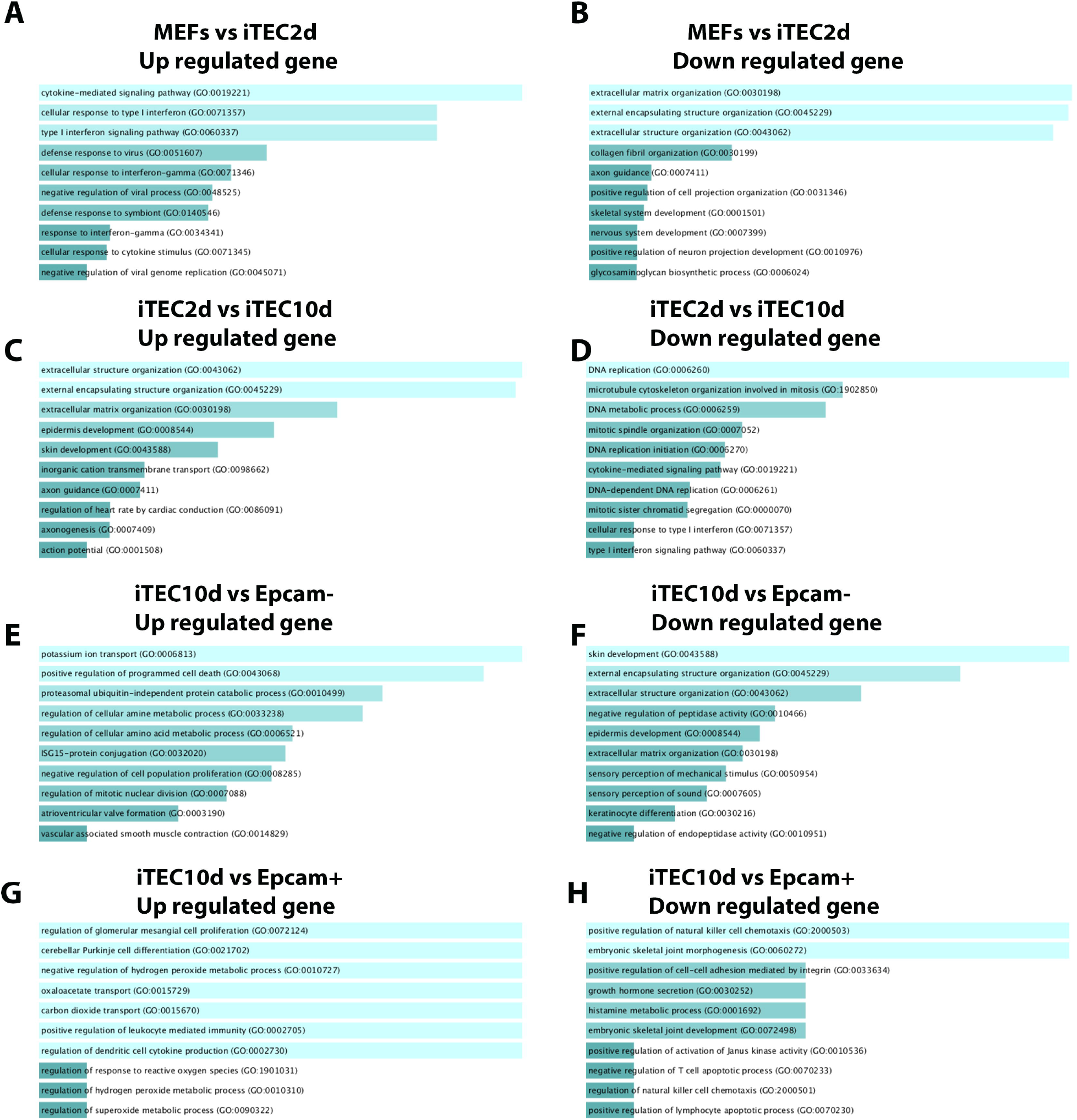
Gene enrichment analysis of up/down regulated genes at different reprogramming stages. A), B) Gene enrichment analysis for MEFs vs iTEC2d up (A) and down (B) regulated genes; iTEC2d vs iTEC10d up (C) and down (D) regulated genes; iTEC10d vs Epcam-up (E) and down (F) regulated genes; iTEC10d vs Epcam+ up (G) and down (H) regulated genes.

Together, our results show that the reprogramming process proceeds through intermediate steps that are significantly different from both MEF controls and the late reprogramming stage. These data indicate that each stage of reprogramming is characterized by a unique gene expression profile, rather than a gradual accumulation of gene expression changes that build through reprogramming.

### Cell cycle arrest is a barrier to the iTEC reprogramming process

The cell cycle arrest seen during iTEC reprogramming is similar to that seen in other reprogramming systems (Treutlein et al., 2016). To investigate this cell cycle arrest at the molecular level, we interrogated our RNA-seq data for changes in cell cycle—associated genes during the early and late stages of reprogramming. Our expression data showed that cell cycle inhibitors including Rb, Cdkn1a/p21, and Gadd45b were upregulated beginning at the iTEC2d stage, while cell cycle-promoting genes such as Myc, Mcm7, and p53 were downregulated in the late stage of reprogramming (Fig4A, B).

**Figure 4:**
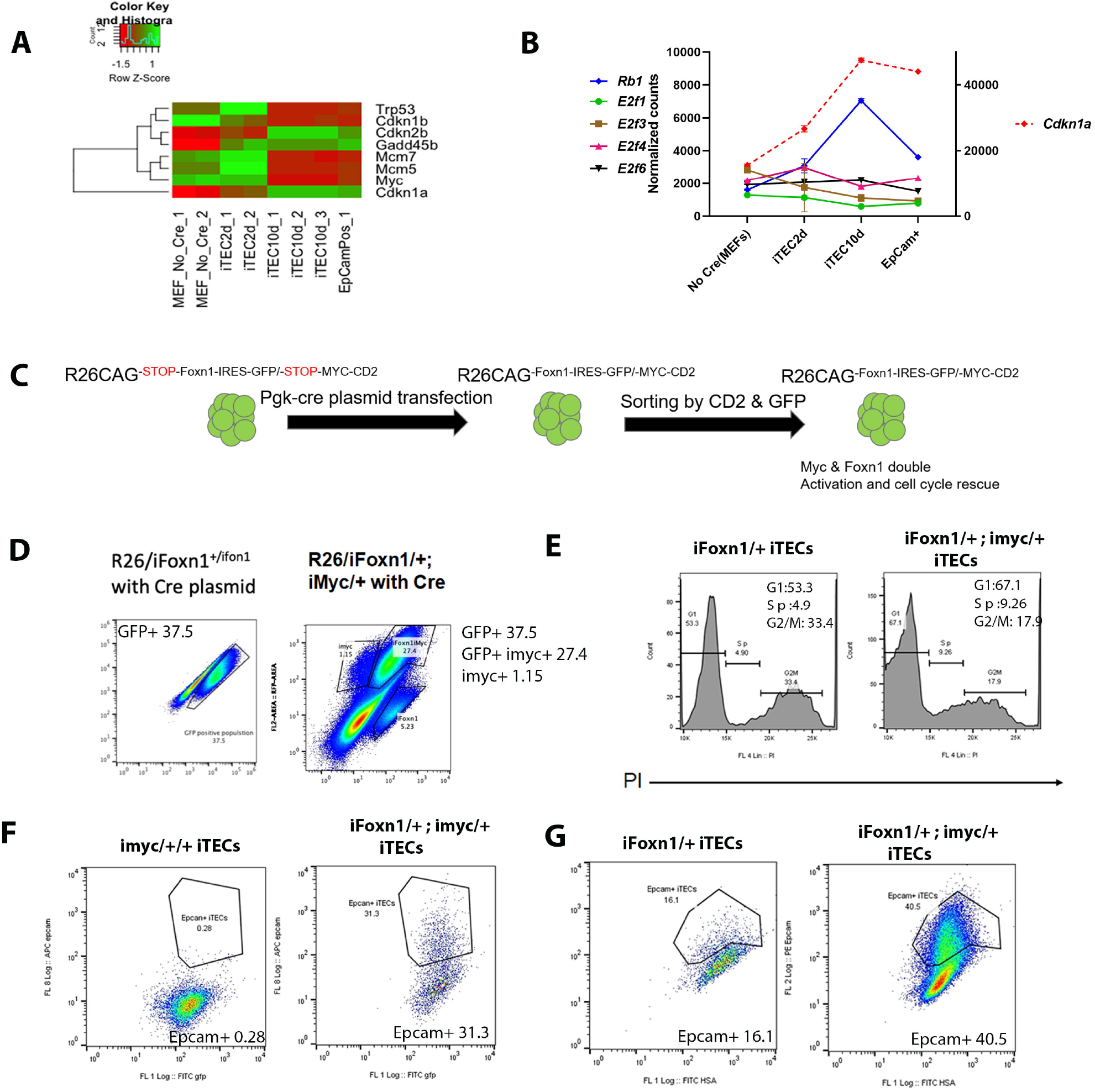
Cell cycle analysis of iTECs and cell cycle rescue using iMYC transgene. A) and B) Gene expression analysis of cell-cycle related genes during iTEC reprogramming. C) Experimental design of inducing iTEC proliferation by overexpressing iMyc. D) Flow cytometric analysis shows that activation of the iMyc transgene leads to about 30% iTECs expressing both iFoxn1 and iMyc. E) PI staining shows significant increase in S phase frequency indicating a rescue of the cell cycle by overexpressing iMyc during iTEC induction. F) EpCam staining showing that iMyc alone was unable to induce reprogramming or EpCam expression. G) Comparison of iFoxn1 alone and iFoxn1 with iMyc shows that Myc expression increases reprogramming efficiency.

*In vivo,* FOXN1 has been shown to promote TEC proliferation (Chojnowski et al., 2014; Nowell et al., 2011). Thus, this cell cycle inhibition is consistent with reprogramming in general, but inconsistent with the known functions of FOXN1. Furthermore, this block results in an inability to expand iTECs in culture, limiting their utility as an experimental system. If cell cycle arrest is necessary for iTEC reprogramming as it is for reprogramming other cell types, then if we rescue the cell cycle, the iTEC reprogramming could be inhibited. Conversely, if cell cycle arrest is not necessary, then the cell cycle’s rescue would not affect iTEC reprogramming and could allow iTEC expansion in culture. To test these possibilities, we utilized an inducible Rosa26 transgene for the Myc gene, which can also be activated by Cre plasmid transfection, and also expresses the human CD2 antigen so that the Myc expressing cell can be detected by staining for huCD2 (Calado et al.). We generated double transgenic MEFs (iFoxn1/+;iMYC/+) in which Cre expression during initial iTEC induction should activate both Foxn1 and Myc gene expression. (Fig. 4C). Cells that have activated both alleles are GFP+ and CD2+. Using PI staining and Cell trace analysis, we find that Myc overexpression can partially rescue the cell cycle arrest in iTECs. iFoxn1+iMyc+ iTEC cultures had increased S phase frequency (Fig. 4D) and a dramatic shift to the left in Cell Trace analysis, indicating dilution of the Cell Trace reagent consistent with most cells undergoing proliferation (Fig. 4E). Activation of Myc alone was unable to induce reprogramming or EpCam expression as detected by flow cytometry (Fig. 4F).

We then analyzed whether this rescue of cell cycle arrest would affect iTEC reprogramming. Surprisingly, activation of the cell cycle does not block iTEC reprogramming, but instead further increases the reprogramming efficiency based on the frequency of induction of EpCam expression. (Fig. 4G). These results suggest that the cell cycle block may be a sideeffect of the reprogramming process rather than a necessary step, and suggests that rescuing the cell cycle arrest can significantly improve the efficiency of iTEC reprogramming.

### cTEC and mTEC genes are activated at different stages of reprogramming

The initial iTEC report showed formation of a well-organized organoid and differentiation of iTECs into both cTEC and mTEC upon transplantation under the kidney capsule (Bredenkamp et al., 2014b). However, it did not characterize cTEC and mTEC lineage markers in detail in iTECs in culture. We performed MA plot analysis to investigate TEC lineage specification and differentiation-related gene expression patterns during the reprogramming process (Fig. 5A-E).

**Figure 5:**
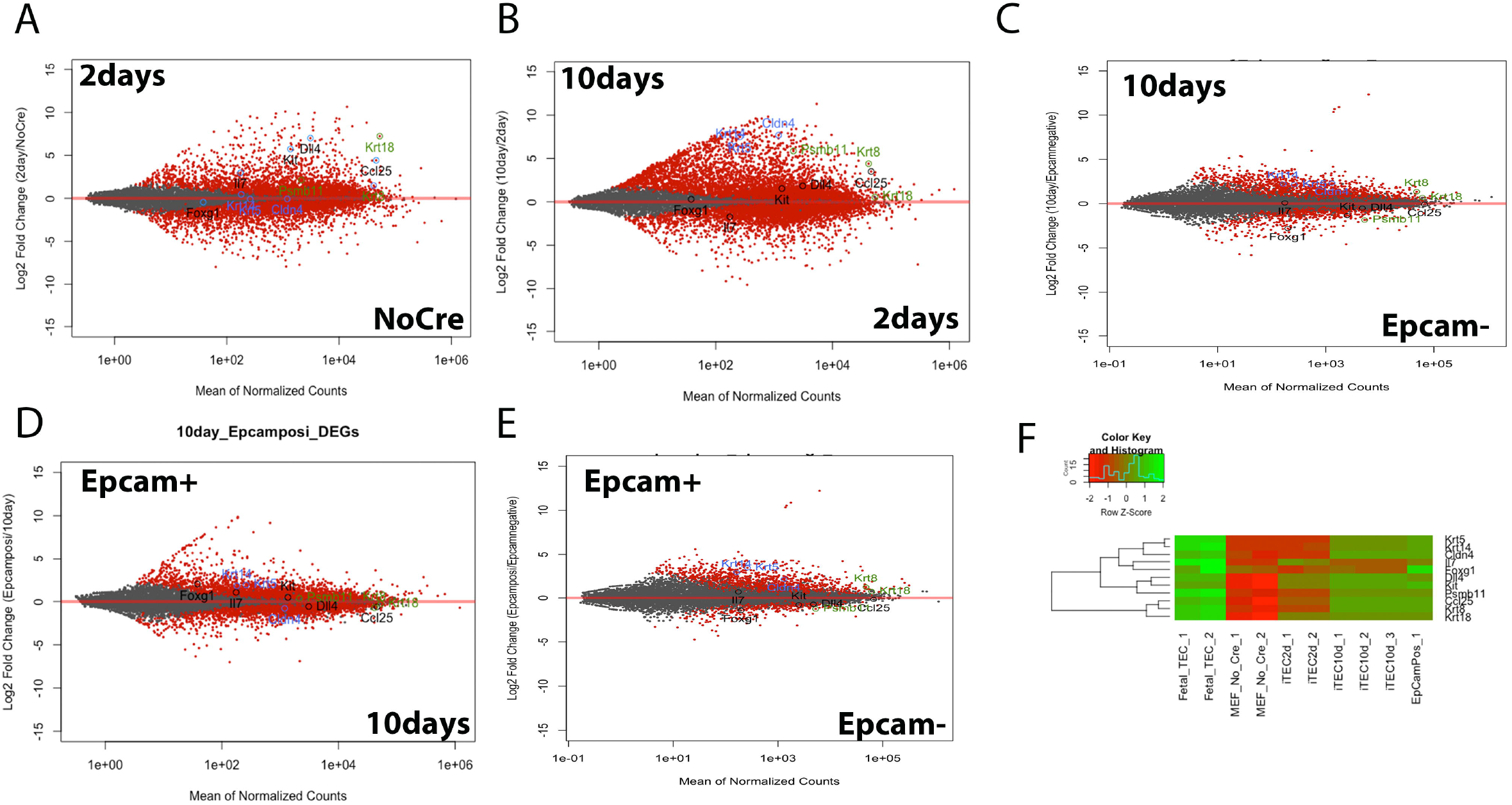
Detailed gene expression comparison analysis of different stage of iTECs. A) – E) MA plot analyses comparing successively more advanced pairs of iTEC reprogramming stages with selected mTECs/cTEC markers highlighted. F) Heatmap analysis of different stages of iTECs reprogramming with important mTECs/cTEC markers. Fetal TEC is also presented in this figure for comparison.

Comparison of MEFs to 2d (Fig. 5A) shows up-regulation of FOXN1 target genes *Dll4, Kit-L*, and *Ccl25*. Strikingly, cTEC markers appear very early, with Krt18 and to a lesser extent Krt8 already significantly upregulated at iTEC2d. Psmb11 (β5t), a cTEC marker and direct Foxn1 target, is also upregulated, although it is expressed at a lower level than Krt8 and Krt18. In contrast, mTEC markers such as Krt5, Krt14, and Cldn4 are not detected at early reprogramming stages (Fig. 5A), indicating that cTEC markers are expressed before mTEC markers during iTEC reprogramming.

Between 2d and 10d, FOXN1 target genes are further upregulated. More remarkable is very strong up-regulation of both cTEC (Krt8 and Psmb11) and mTEC markers (Fig. 5B). mTEC markers first appear at iTEC10d stage, where they are highly differentially expressed compared to 2d samples, but still at much lower expression levels than the cTEC markers. Interestingly, while Foxn1 target genes and cTEC markers remain stable from the iTEC10d to EpCam+ iTEC (Fig. 5D), the mTEC related genes Krt14 and Krt5 are differentially upregulated at this latest stage of reprogramming. (Fig. 5E). Notably, mTEC but not cTEC markers are also upregulated in EpCam+ iTECs compared to EpCam-iTECs. This progressive up-regulation of TEC differentiation genes with delayed mTEC marker up-regulation is further evident in a heat map analysis (Fig. 5F). However, EpCam+ iTEC have much lower expression of these markers than fetal TEC. This comparison is more fully explored below.

### Notch signaling inhibits mTEC differentiation during iTEC reprogramming

The differential lineage-specific gene expression analysis showed that the mTEC progenitor marker Cldn4 is one of the highest expressed mTEC markers at later iTEC stages, while the terminal differentiation marker Aire is not detected in our RNA-seq data (Fig. 5). Thus, we hypothesized that mTEC differentiation is induced at the progenitor stage during reprogramming, but that further mTEC differentiation is inhibited or incomplete. Recently we and others showed that the Notch pathway plays an essential role in mTEC proliferation and differentiation (Li et al., 2020; Liu et al., 2020). Most critically, while Notch signaling is required to specify mTEC progenitors and their initial proliferation, too much Notch signaling after initial mTEC specification would lead to a block of mTEC terminal differentiation, in which mTECs would arrest at Cldn4+ mTEC progenitor states. These *in vivo* mechanisms are similar to what we observe in iTEC reprogramming and suggest the possibility that persistent Notch pathway may suppress mTEC differentiation during iTEC reprogramming, similar to how it functions in mTEC differentiation *in vivo*.

To test this hypothesis, we first analyzed gene expression of components of the Notch signaling pathway. (Fig. 6A, B). We found that multiple Notch receptors, ligands, and downstream target genes were upregulated during reprogramming and then maintained throughout the late reprogramming process. These data indicate the induction and persistence of Notch signaling during reprogramming. To test whether this maintenance of Notch signaling is the reason why mTEC differentiation is blocked at a Cldn4+ stage, we designed a temporally controlled Notch inhibition experiment (Fig. 6C). We allowed early reprogramming to progress in order to establish initial mTEC identity, then at day 5 after the initial transfection added the Notch signaling inhibitor DAPT daily, culturing the cells for a further 7 days as usual. DAPT-treated cultures reduced but did not eliminate Notch signaling as indicated by a 50% reduction in Hes1 expression (Fig. 6D). Cldn4 was also downregulated to a similar degree, indicating that iTECs had reduced mTEC progenitor characteristics under DAPT treatment. To assess mTEC differentiation, we used *Aire* expression, which is a critical marker for mTEC maturation and terminal differentiation. Aire expression was detected in 6 out of 11 experiments, and in 5/6 was significantly upregulated in the DAPT cultures compared to DMSO controls. These data suggest that inhibiting the Notch signaling pathway during later stages of reprogramming allowed mTEC lineage differentiation, as predicted by the *in vivo* role for Notch signaling in mTEC differentiation. As an independent assessment of mTEC differentiation, we also measured Fezf2, an Aire-independent marker of mTEC differentiation Fezf2 was not consistently different between DMSO and DAPT-treated cultures. Inhibiting the Notch signaling pathway also did not affect iTEC reprogramming efficiency based on the percentage of the EpCam+ population and did not induce MHCII expression (Fig. 6E).

**Figure 6:**
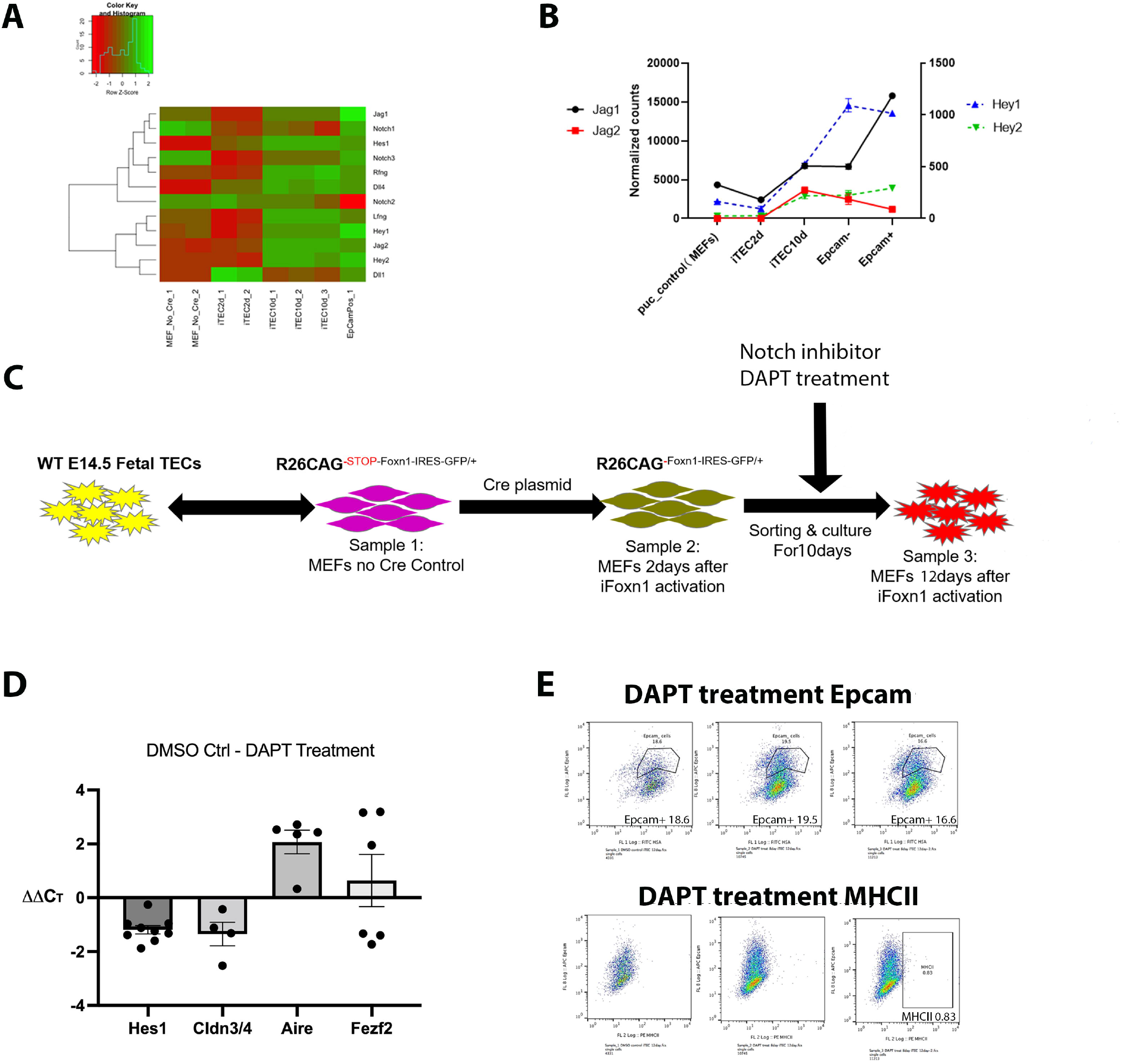
Notch pathway analysis and DAPT treatment experiment. A) and B) Gene expression heatmap & line plot of components of the Notch signaling pathway. C) Experimental design of DAPT treatment experiments for activating Notch signaling. D) qRT-PCR analysis of DAPT treatment effect on Notch pathway target gene Hes1 and mTEC differentiation related genes Cldn3/4, Aire, and Fezf2. E) Epcam/MHCII flow cytometry analysis of DAPT treated iTECs

Taken together, these data suggest that the upregulation of *Aire* was a specific response to inhibiting Notch signaling during iTEC reprogramming. These data indicate that iTEC reprogramming may follow the same mTEC differentiation mechanisms *in vivo* and opens the possibility of establishing iTEC as a useful system for investigating mTEC differentiation and function.

### EpCam+ iTECs are missing specific genetic signatures found in fetal TECs

To test how similar iTECs in culture are compared to *in vivo*-derived fetal TECs at the transcriptome level, we isolated and sequenced total fetal TECs at E14.5 as well as a second sample of EpCam+ iTECs and analyzed these data in combination with out previously collected samples. PCA analysis clearly demonstrates that all iTEC samples are still quite different from fetal TECs, although the Epcam+ samples are the closest (Fig. 7A). The sample distance count matrix shows a similar result, with fetal TECs distinct from all iTEC samples, and Epcam+ samples as the closest to fetal TECs. (Fig. 7B). GO term analysis indicated that fetal TEC overall have higher expression of cell adhesion molecules relevant to epithelial identity and TEC differentiation (including Claudins, CD40, and CD28) and a variety of immune function related gene categories (Fig. 7C), while EpCam+ iTECs have higher expression of a variety of categories indicating retention of mesenchymal and/or fibroblast-like characteristics (Fig. 7D). MA Plot analysis of EpCam+ iTEC vs fetal TEC also showed that both mTEC and cTEC markers were higher expressed in fetal TEC, although the degree of difference was less in 10d vs 2d TEC (Fig. 7E, F). While direct FOXN1 targets such as Dll4, Ccl25, and Kitl were significantly higher in fetal TEC compared to iTEC2d, by the EpCam+ iTEC stage FOXN1 targets were similar in level to fetal TEC, suggesting that the differences between iTEC and fetal TEC are not totally driven by differences in *Foxn1* expression levels.

**Figure 7:**
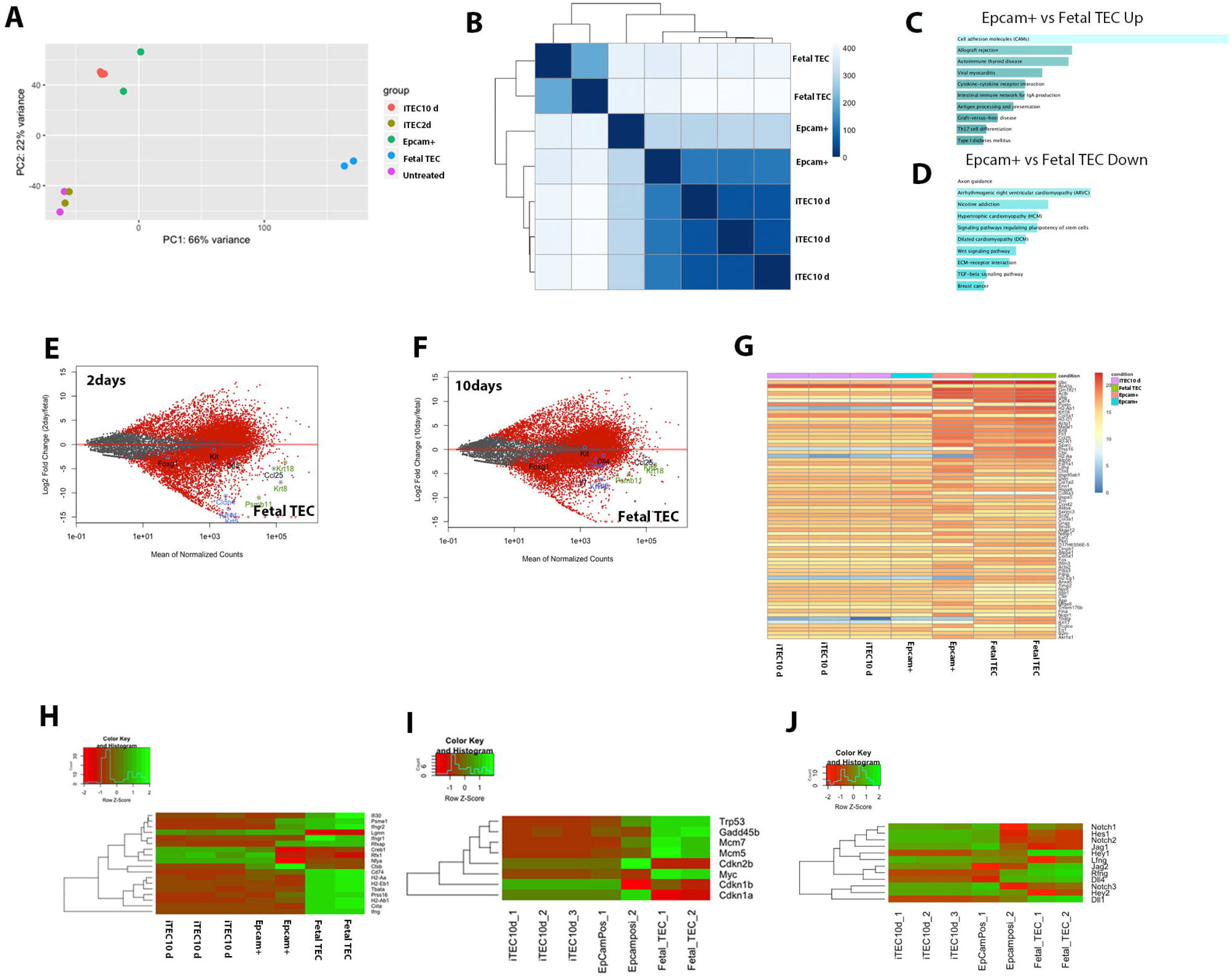
Transcriptome and differential gene expression analysis of comparison between iTEC and E14.5 Fetal TEC shows EpCam+ iTECs are missing specific genetic signatures. A) PCA analysis of different time points of iTEC samples and fetal TECs. B) Sample distance analysis of iTEC 10day, Epcam+ iTEC, and fetal TECs. C, D) Enrichment analysis for Epcam+ iTEC vs Fetal TEC up & down regulated genes. E, F) Gene enrichment analysis comparing 2days and 10days iTEC stages to fetal TECs for seected mTEC and cTEC markers. G) Top differentially expressed genes between different iTEC reprogramming stages and fetal TECs. H – J) Heatmap analysis of top differentially expressed genes between iTEC reprogramming and fetal TECs by category: H) MHCII related genes, I) cell cycle related genes, and J Notch pathway related genes.

We then investigated the gene expression differences between iTEC and fetal TEC in more detail. We first used a count matrix heatmap to identify differentially expressed genes that define these different stages (Fig. 7G). This analysis clearly shows that the two EpCam+ samples are not just different from each other, but that one is clearly more similar to fetal TECs, indicating that it represents a more advanced stage of reprogramming. Four of the genes that fail to be up-regulated in any iTEC samples compared to fetal TEC (blue/white in all iTEC samples, red/orange in fetal TEC) are three MHCII genes (annotated as H2-A or H2-E) and Tbata, which is known to regulate TEC proliferation *in vivo* (Flomerfelt et al., 2010).The gene that is the most downregulated in fetal TEC compared to all iTEC samples is Col6a3 (white in fetal TEC, orange in iTEC), consistent with persistent retention of some fibroblast characteristics in iTECs. Heatmap analysis of a smaller subset of 18 of the most highly DEGs (Fig. 7H) further highlighted the conclusion that lack of MHCII-related genes was a defining difference between iTECs and fetal TEC, with 10/18 of these DEGs genes associated with MHCII, including Ciita, the major transcriptional regulator of MHCII expression (León Machado and Steimle, 2021). In addition, 3 genes are associated with TEC function and/or proliferation (*Psem1, Prss16, Tbata*) and one with Notch signal modulation *(lfng).* This heatmap analysis also demonstrated that the more differentiated EpCam+ sample did share some of these characteristics with fetal TEC, including up regulation of some MHCII-related genes (Psme1, Rfxap) and Ifngr1, and down regulation of Creb1, Rfx1, Nfya, and Ctsb similarly to fetal TEC. Similar patterns were seen with cell cycle related genes (Fig. 7I) and Notch pathway related genes (Fig. 7J); in both cases one of the two EpCam+ samples were more similar to fetal TEC, although still retaining significant differences. These data are consistent with our results above, and suggest that these pathways may be potential candidates to target for improving iTEC protocols to be more similar to fetal TEC.

To sum up, by analyzing the transcriptome of fetal TECs, we find that even the latest stage of iTEC analyzed, EpCam+ iTECs, are distinct in gene expression profiles from fetal TECs. However, Epcam+ samples share more similarities with fetal TECs than earlier stages. A critical difference is the expression of MHC Class II genes, which are still absent in iTECs, but are critical functional components of TECs.

## Discussion

The results of this study provide a significant advance in our understanding of iTEC reprogramming and provide clear avenues for future improvement of this technique, although many questions need further study. We have shown that iTEC reprogramming is not linear, and defined characteristics of early and late stage reprogramming. Our data indicate that reprogramming begins with immediate up-regulation of FOXN1 direct target genes that are not normally expressed in MEFs, consistent with the role of Forkhead transcription factors as pioneer factors (Zaret and Mango, 2016). Reprogramming then expands to include many additional changes not directly attributable to known FOXN1 *in vivo* functions. We further show that the cell cycle arrest that occurs early in iTEC reprogramming is not required for reprogramming to occur, and instead inhibits reprogramming efficiency. Further, we show that iTEC reprogramming follows a path consistent with the “cTEC first” model, with mTEC markers appearing later, and consistent with arrest at the mTEC progenitor stage. Finally, we show that this apparent block to mTEC differentiation can be partially relieved by inhibiting the Notch signaling pathway, suggesting that mTEC differentiation in iTECs follows similar pathways as those of endogenous fetal mTECs. Finally, we identify the MHC pathway as a major remaining difference between iTEC and fetal TEC.

One persistent question is why iTEC differentiation is not uniform across iTEC cultures. In our current iTEC direct reprogramming protocol, the EpCam positive population is always a subset of the cultures, even though all iTECs are expressing the same Foxn1 transgene for the same amount of time. One possibility is that the cell cycle may play a partial role in this process, since rescuing the cell cycle block did increase the percentage of Epcam+ iTECs (although it still was not 100%). MEFs are a heterogeneous population, and it is possible some are more susceptible to FOXN1-reprogramming than others (Shakiba et al.). Another variable is endogenous *Foxn1* expression. We consistently detect activation at the endogenous locus at a low-level at the late stage of reprograming, although it is unclear whether this occurs in all cells, or whether this activation is somehow involved in reprogramming. Additional analyses of heterogeneity during reprogramming needs to be done to test these hypotheses.

Several previous studies show that cell cycle arrest is necessary for direct lineage reprogramming of other cell types, such as neuron direct reprograming (Treutlein et al., 2016). However, our data show that while cell cycle arrest occurs, it is not necessary for iTEC direct linage reprogramming. In contrast, removing this cell cycle block by forced Myc expression resulted in higher efficiency of EpCam+ iTEC generation. However, it is still unclear what exactly cell cycle does in the iTEC reprogramming process, and to what extent these proliferating iTECs are similar to or different from cell cycle arrested iTECs. It is also possible that Myc overexpression has more impacts on iTEC reprogramming than on cell cycle regulation, as Myc both alone and as a component of the RB pathway is itself implicated in TEC differentiation (Cowan et al., 2019; Garfin et al., 2013).. However, Myc expression alone had no detected impact in the absence of Foxn1 expression. Further experiments will need to be done to detailed analyze the impact of cell cycle arrest rescue and further understand the mechanisms underlying how cell cycle interacts with iTEC reprogramming.

Finally, we do detect both cTEC and mTEC marker expression during iTEC reprogramming. Our data are consistent with both the “cTEC first” model of TEC differentiation (Takahama et al., 2017) and with successful induction of mTEC lineage specification, at least to the Cldn4+ mTEC progenitor stage. Our experiments also demonstrate that persistent Notch signaling may limit mTEC differentiation in iTEC cultures, and that blocking Notch signaling at later stages can result in significant mTEC differentiation, including expression of the critical marker *Aire.* However, it is still unknown how this process is occurring at the individual cell level. It is possible that all iTECs expresses cTEC markers first, and as they further differentiate during the reprogramming process, all or a subset start to express mTECs markers. Alternatively, it is possible that different subsets of cells initiate cTEC and mTEC differentiation directly. Resolution of this question will require analysis of reprogramming at the single cell level.

Finally, none of our iTEC cultures express MHC Class II, a crucial functional hallmark of TEC differentiation. In the original iTEC report, crosstalk with immature thymocytes either in culture or after transplant induced MHC Class II expression, allowing for the generation of single positive CD4+ T cells(Bredenkamp et al., 2014b). Thus, future experiments to identify the crosstalk elements that activate MHC Class II expression will be essential to generation of a fully viable iTEC culture system as a novel experimental platform for understanding TEC biology and function, and as a potential approach for the eventual generation of human iTEC for *in vitro* or *in vivo* therapeutic uses.

## Materials and Methods

### Mice

Rosa26^CAG-STOP-Foxn1-IRES-GFP/+^ mice (Bredenkamp et al., 2014b) were backcrossed onto the C57BL/6J genetic background for at least 5 generations then maintained by intercrossing. Each batch of mouse embryonic fibroblasts (iFoxn1+ MEFs) were pooled from embryos in a single litter of E13.5 embryos generated from a Rosa26^CAG-STOP-Foxn1-IRES-GFP/+^ female crossed to a C57Bl/6J male. For timed matings, noon of the day of the vaginal plug was taken as day 0.5.

### MEF isolation

MEFs were prepared from E13.5 embryos decapitated and stripped of all internal organs and trypsinized into a single-cell suspension. Cells were plated in DMEM containing 10% fetal calf serum, 2 mM sodium pyruvate, 4 mM glutamine, 50 μg/ml streptomycin and 50 U/ml penicillin (DMEM/FCS). Each embryo was genotyped using the following primers to detect the iFoxn1 allele: iFoxn1 Forward Primer 5’ – TGG AGT AGG CGG GGA GAA GG −3’ iFoxn1 forward Primer Mid: 5’—TCG CCC TTC CCA ACA GTT GC −3’ iFoxn1 Reverse Primer: 5’ −GCC CAC ACA CCA GGT TAG CC—3’. MEFs were freshly isolated and initially cultured in 60mm plates until reaching 80% confluency, and transferred to T-75 flasks until reaching 80~90% confluency. Then, MEFs were trypsinised and frozen in −80’C overnight. Cryovials of MEFs were then transferred into liquid nitrogen tanks until usage. Once MEFs were thawed, they were expanded once or twice more for downstream experiments.

### Cell Culture & Transfection

Primary iFoxn1/+ MEFs were thawed then cultured in DMEM containing 15% fetal calf serum, 2 mM sodium pyruvate, 4 mM glutamine, 50 μg/ml streptomycin and 50 U/ml penicillin (DMEM/FCS). Cells were passaged once at 70% – 80% confluency, then collected at 70% confluency for Nucleofection. Primary iFoxn1/+ MEFs for Nucleofection experiment were passaged no more than than 5 times for all analysis in this research. Nucleofection experiments were performed following the manufactures’ protocol for mouse embryonic fibroblasts (Amaxa™ Nucleofector™, Lonza). To induce iFoxn1 expression, pPGK-Cre-bpA (Addgene # 11543) was used in the transfection according to the manufacturer’s protocol. After Nucleofection, cells was further cultured for 48 hours until sorted by flow cytometry.

### Flow cytometry & Sorting

Adherent MEFs or iTECs were collected by trypsinization. For fetal TEC collection, E15.5 fetal stage thymi were dissected and digested in 1 mg/ml collagenase/dispase (Roche, Basel, Switzerland) and passed through a 100 μm mesh to remove debris. Thymi were processed individually before genotyping, then pooled. PE-Cy7 conjugated anti-CD45 (BioLegend, 30-F11, 1:150) and APC-conjugated anti-EpCam (BioLegend, G8.8, 1:150) were used to isolate TEC populations. Anti-MHCII (BioLegend, 1:150) was also used for TEC analysis. Cell sorting was performed using a BioRad-S3 cell sorter.

### Notch Signal Inhibition Experiment

Exactly 60 hours (5 days) after transfection, 150,000 iTECs were treated with No Treatment, DMSO control, or 10μM of *γ*-secretase inhibitor, DAPT (GSI-IX) (Apex Bio, Catalogue A8200). The treatments were continued once every 24 hours for 7 consecutive days. Two or three technical replicates and at least 5 biological replicates were done for analyzing the effect of Notch signal inhibition on iTECs. Cells were then collected as below for RNA extraction.

### RNA extraction, Reverse transcription, and quantitative PCR (RT-qPCR)

RNA was prepared using the RNAeasy mini kit (Qiagen) according to the manufacturer’s instructions. All samples were DNase treated during the preparation. cDNA was synthesized using the iScript Reverse Transciption Supermix for RT-qPCR (Bio-Rad), according to the manufacturer’s instructions. Thermal cycling conditions for the cDNA synthesis were as follows: 25°C for 5 minutes; 46°C for 20 minutes; 95°C for 1 minute. Quantitative PCR was performed with Applied BiosystemsTM 7500 Real-Time PCR System (ThermoFisher Scientific). Thermal cycling conditions were as follows: 50°C for 2 minutes; 95°C for 10 minutes; 40 cycles of 95°C for 15 seconds, 60°C for 1 minute. Relative expressions were determined after normalization to the geometric mean of two reference genes (Gapdh and Tbp), and were determined after normalization to Gapdh whenever Tbp was undetected. All samples were run with two or three technical triplicates and with no RT controls. The primers used for RT-qPCR are purchased from ThermoFisher Scientic, TaqMan Gene Expression Assays. All probes were designed with FAM-MGB Reporter-Quencher.

### Bulk RNA sequencing & downstream processing, analysis and graphic display of RNA-seq Data

Bulk RNA sequencing was performed in Georgia genomics and bioinformatics core (GGBC) using an Illumina NextSeq 500 sequencing platform to generate single end sequencing data. Raw reads were pre-processed with sequencing grooming tools FASTQC, Trimmomatic and then further assembled by Hisat2, SAMtools, and StringTie. Differential gene expression analysis between samples and graphical display (including count matrix plot, PCA, heatmap and other plots) were performed using DEseq2 and R package ggplot2. Gene ontology enrichment analysis was performed using online EnrichR program (Chen et al.; Kuleshov et al.).

## Acknowledgements

We thank Clare Blackburn for helpful advice and discussions during the progress of this project. We thank J. Nelson in the Center for Tropical and Emerging Global Diseases Flow Cytometry Facility at the University of Georgia for flow cytometry and cell sorting technical support. This work was supported by the Biomedical Microscopy Core and the Georgia Genomics and Bioinformatics Core facilities.

## Author contributions

Conceptualization: Z.M, B.C., N.R.M.; Validation: S.K., Z.M.; Formal analysis: Z.M., S.K., B.C., N.R.M.; Writing – original draft: Z.M., N.R.M.; Writing – review & editing: Z.M, S.K., N.R.M.; Supervision: B.C., N.R.M.; Project administration: N.R.M.; Funding acquisition: N.R.M.

